# TickMotion: A user-friendly tool to measure and visualize muscle contractions and movements in insect and other arthropod tissues

**DOI:** 10.1101/2025.07.05.662616

**Authors:** Matej Medla, Dušan Žitňan

## Abstract

TickMotion is a Python-based software tool designed for the post hoc analysis of video recordings that capture movements or muscle contractions in insects. It provides a simple, fast, and automated approach for semi-quantitative assessment of motion by measuring the “pixel activity” – changes in pixel intensity or color caused by movement. Unlike many existing approaches, it does not require specialized equipment, chemical reagents, or expensive software. In this context, clusters of pixels represent biological structures such as tissues or entire organisms, with their activity reflecting dynamic changes during the recording. The application can distinguish active pixels from inactive ones within selected regions of interest (ROIs) and measure their intensity over a pre-set time. It offers two distinct analyses: Graphical Analysis (GA), and Pixel Activity Analysis (PAA), both customizable to accommodate various experiments. GA generates a graphical illustration of pixel activity (in selected ROIs) over time together with numerical data from the measurement. PAA produces PNG images highlighting the active regions (clusters of pixels) in the video. Initially designed to analyze muscle contractions in tick tissues, TickMotion is useful for applications extending to any movement analysis. It serves as a simple but valuable tool across diverse research areas.

## INTRODUCTION

Python is one of the most robust and widely used programming language, and serves as an invaluable tool for various analyses. In the muscle contractions analysis, there is only a handful of online applications, such as EthoVision XT, DeepLabCut, and ANY-Maze (Bailoo et al., 2010; Mathis et al., 2018; Bühler et al., 2023). Although these tools are the similar to TickMotion, they originally were designed for distinct purposes, most commonly movement and behavior tracking in small mammals.

In general, muscle contractions and movement analysis in arthropods or their tissues can be performed by various techniques, the most common is electromyography (EMG) measurement. Fully-established in insects in the mid-20th century, EMG remains widely utilized in many laboratories due to its ability to provide detailed and precise measurements of muscle contraction levels across diverse organisms (Yoshida et al., 2023; Hermanson and Altenbach, 1981; Vo-Doan et al., 2022; Pflüger et al., 2011). However, utilizing this technique requires specialized equipment and computational tools for data processing. Additionally, while EMG excels in quantifying muscle activity, it does not provide visual representation of muscle contractions.

Our approach was to establish a new, simple method to analyze movements and muscle contractions within various tissues or entire animal using a post-experimental video analysis. In this study, we introduce TickMotion that was engineered to quantify “pixels activity”. The application converts the input video into grayscale, which simplifies the visual data for subsequent analysis. Every recorded moving particle can be seen as a group of pixels whose colors or shades change over time, while the color and shade of pixels in the (stationary) background remains the same. The application can automatically identify “active pixels” (those that exhibit changes in color and/or shade) in a video and label them as white (or red, depending on the selected script), while inactive pixels are labeled black. This allows for selective semi-quantitative measurement of the intensity of color and shade changes of pixels in the video, over a defined period. TickMotion offers 2 different analyses: (i) Graphical Analysis (GA), and (ii) Pixel Activity Analysis (PAA). Additionally, our tool is open-sourced, deposited on GitHub repository and includes two versions of the scripts: TickMotion and TickMotion v_2, with the latter one featuring a modified PAA. TickMotion v_2 shows active pixels as red, while the background is derived from the first frame of the video file.

Arthropods, the most abundant and diverse group of organisms on Earth play important roles in the ecosystem, ranging from essential commensals to parasitic species. Understanding their way of life is crucial for developing strategies to either ensure their protection or mitigate the transmission of pathogen-causing diseases by reducing their development, physiology or reproductive fitness. Most studies are focused on biological functions, such as feeding, growth, ecdysis, reproduction, locomotion, and behavior. These studies usually include video recordings to observe and track various behavior and to identify differences between control and test groups. However, post-experimental analysis of such recordings can be labor-intensive and is often limited to qualitative observations or general descriptions. In many cases, the need for objective, quantitative data is critical – especially when assessing subtle physiological or behavioral changes.

Additionally, numerous bioactive molecules are known to affect arthropod physiology by stimulating or inhibiting muscle contractions of various tissues during feeding, ecdysis, excretion, and reproduction (Šimo and Park, 2014; Hewes et al., 1998; Hu et al., 2018; Onken et al., 2004; Matsumoto et al., 2019; Kim et al., 2015). In this context, TickMotion emerges as a user-friendly, precise, sensitive, and efficient tool for detection of specific movements during these processes in a video recordings. It offers customization options to filter background noise, measure only specific parts of the video, and provides all features directly within the application. Moreover, its simple and practical interface rapidly delivers valuable results, making it applicable in various research fields.

## MATERIALS AND METHODS

### Experimental animals

All animal experiments were carried out in accordance with the Animal Use Protocol approved by the ethical committee of Biomedical Research Centre of Slovak Academy of Sciences and the State Veterinary and Food Administration of the Slovak Republic (facility number SK U CH01016, permit number 3954/19-221 and 292/16-221k). *I. ricinus* ticks from the breeding facility at the Institute of Zoology SAS were kept in tubes in 95% humidity chambers under a 10:14 h light/dark cycle at 22°C. Adult ticks were fed on rabbits in rubber chambers glued onto the shaved backs of the animals covered with a fine mesh net. Females were placed in the feeding chambers with males in 1:1 ratio, as described (Jones et al., 1988; Almazán et al., 2018). Partially-fed females were collected daily between days 6 and 10.

### *In vitro* and *in vivo* functional assays

Ticks at specific stages were dissected in physiological saline (140 mM NaCl, 5 mM KCl, 5 mM CaCl_2_, 4 mM NaHCO_3_, 1 mM MgCl_2_, 5 mM HEPES, pH 7.2), immobilized on a Petri’s dish using a soft dental wax, and placed in the drop of Hank’s solution (Sigma-Aldrich, St. Louis, MO, USA), to prevent desiccation. All experiments were monitored under a binocular microscope Leica M205 C or Leica S9D, and photographed and recorded using a camera Leica Flexacam C1.

The *in vitro* contraction assays were performed on male accessory glands or the female midguts to test the sensitivity of TickMotion in measuring differences between control and treated tissues. For each individual, muscle contractions were measured at three experimental conditions: 10 minutes after the ticks were placed on a Petri dish (control), after the muscle stimulation (stimulated), and following the muscle inhibition (inhibited). The recording time was set to 1-2 minutes.

The Pixel Activity Analysis (PAA) – specifically the one in the second version of the script (TickMotion v_2) is suitable for monitoring *in vivo* locomotion in ticks (see below). Male ticks were individually placed into larger Petri dishes and human breath was used to “activate” them (Vassallo and Pérez, 2002). Recording started immediately after activation and continued for up to 30 seconds.

### Data acquisition and analysis

As the analysis is based on the quality of a video, a stabilized camera directly mounted on the microscope with consistent zoom and focusing area were used for all recordings throughout the video acquisition process. The LED ring light (e.g., Leica LED5000 HDI) was the most suitable to improve the quality of the recording. The protocol is divided into two parts, as the TickMotion offers 2 different analyses: (i) Graphical analysis (GA), and (ii) Pixel activity analysis (PAA). A single video can be analyzed with both, GA and PAA, which complement each other to provide more detailed and comprehensive results. For optimal performance and uninterrupted analysis, it is recommended to follow the steps in each protocol in a top-to-bottom sequence.

Using Graphical Analysis (GA), ROIs were selected within the most active part of male accessory gland (*vas deferens*), with frame parameters: −x = 50, +x = 50, −y = 30, +y = 30 pixels (Suppl. Fig. S1A-D). To compare different responses of the accessory glands across three experimental conditions, the interquartile range (Q3-Q1; representing the middle 50% of measured values, Suppl. Fig. S1F) was used to minimize the impact of extreme values (Whitley and Ball, 2002). The results were presented as percentage amplitude, while stimulated accessory glands set as the maximum at 100%. Additionally, detected contraction levels within selected ROIs were used to create a progress curve reflecting color intensity and shade changes of pixels (y-axis) over a pre-set time (x-axis).

The PAA was used for the visualization of *in vitro* muscle contraction level of the male accessory glands and female midguts, as well as to analyze *in vivo* locomotion of male ticks. The second version of the script that contains modified PAA (TickMotion v_2) was also tested using *in vitro* and *in vivo* approaches.

For an accurate GA and PAA, the experiments were performed on multiple ticks and data were further processed in Excel (Microsoft, Redmond, WA, USA), GraphPad Prism 5 (GraphPad Software Inc.), and Adobe Photoshop CS6 version 13.0 (64bit) (Adobe Systems, San Jose, CA). Video recordings were edited (cropped and trimmed) using Microsoft Clipchamp (Microsoft, Redmond, WA, USA).

### Code availability and installation

The TickMotion code is fully open-source, licensed under the GNU GPLv3 license, and available on GitHub at: https://github.com/medli2568/TickMotion. The repository contains two script versions: “TickMotion” and “TickMotion v_2”, which differ only in the Pixel Activity Analysis (PAA) output. Both scripts can be run directly in a Python Integrated Development Environment (IDE), such as PyCharm, or converted into standalone applications using PyInstaller. PyInstaller packages the Python script into an executable (.exe) file in a few simple steps (see “Code Install” section on GitHub for full instructions). This allows users to run the software without requiring a separate Python installation.

### Graphical analysis (GA)

The GA begins by clicking on the button: *“Graphical Analysis”*. The final outcome of the GA includes a graph showing the amplitude of contractions over time (frames) in the selected ROIs, *txt* files containing data of each ROI featured in the graph, and PNG images representing each analyzed segments of the video file.

In the first step (*“Number of ROI”*), type the number 1 to 10 of ROIs. Each ROI will undergo analysis separately. TickMotion does not support the analysis of more than 10 ROIs simultaneously. Next, click on *“Choose Video and ROI”* to navigate TickMotion to the appropriate folder and select a video file (supported formats: .mp4, .avi, and .mov) by double-clicking. Continue in this step and select (left-click) the ROIs. The application will automatically stop the selecting process and the video file will be closed once the number of selected ROIs corresponds the number established in the previous step. Do not close the video file during selection process. If you intend to change selected ROIs, let the program close it automatically (explained above) and repeat this step directly in the application. The graph shows results from any movement captured on the video. Try to select all ROIs from parts of the video, where movements are clear, muscle contractions are repetitive and/or the background noise is the lowest.

To enhance the efficiency of the analysis, and to target specifically contracting or moving segments of a video file, square frames are generated around each selected ROI (Suppl. Fig. S1A). The frame size is customizable in the step *“Frame Parameters (dimensions)”*, with the default dimensions set as follows (in pixels): “-x” = 50, “+x” = 50, “+y” = 50, “-y” = 50. Changes to frame dimensions must be confirmed by clicking the *“Update Frame Parameters”* button. For optimal muscle contraction analysis, it is essential to select a frame size that closely surrounds the area of interest to minimize background interference.

Right after clicking on the *“Analyze”* button, the graph is automatically generated by measuring pixel activity within the selected ROIs and specified frame dimensions from the previous steps. This graph incorporates data for all ROIs, with the video file duration set to a predetermined 10 minute length (maximum). Data for each ROI in the graph are normalized based on the pixel intensity of the first frame, ensuring that the initial x value for every ROI is always 0, in the final graph. The system automatically saves all data in the designated “Output Folder”. PNG images are derived from the initial video frame and aim to facilitate the video analysis during the setup of frame dimensions in preceding steps. The whole analysis took around 30 seconds per 1 minute long video (recorded at 60 fps) and 1 ROI selected. Once the analysis is completed, you can revise the frame dimensions around selected ROI and repeat the analysis, if needed.

TickMotion also offer possibilities to easily enter the Output Folder, return back within the application and terminate the application. Note that files from one analysis will be named corresponding to each ROI separately. The output files from multiple analyses will always have the same name, so it is necessary to save the data to different folders.

### Pixel Activity Analysis (PAA)

The PAA starts by clicking on the button: *“Pixel Activity Analysis”*. As a result, TickMotion provides a PNG image of active and inactive pixels from the video. It calculates the activity of each pixel separately, when it met the preset settings, active pixels are shown as white and inactive as black. The TickMotion repository on GitHub also includes the second version of the script, named “TickMotion v_2”, where the PAA is modified by visualizing the results (not the method). This approach was designed to analyze movements on a bigger scale.

In the first step, click on *“Choose Video”* button in the PAA, navigate to the folder, and select a video file (again, multiple video formats are supported). ROIs selection is not needed in this step, as the PNG are generated for the whole video automatically. Similarly to GA, click on the *“Select Output Folder”* button and navigate to the appropriate folder, where the PNG image will be saved. The output PNG are named “active_pixels” in every analysis. Therefore multiple analyses have to be saved in different folders, or renamed after analysis manually.

Subsequently, the application allows to customize a video analysis by selecting “Threshold percentage” (t%) and “Frame skip” (FS) parameters. T% refers to the activity of pixels per time and is preset to 80. Setting t% to 80 means that every pixel that is active (changes its color/shade) for at least 80% of video duration appears white, otherwise it appears black (depending on the script). This feature allows you to reduce the background bias to minimum by setting higher t%. Other parameter – FS refers to the number of frames that will be analyzed. The FS parameter is set to 0 by default, meaning no frames are skipped and every video frame is analyzed. However, for extended recordings captured at 60 fps (optional), it is recommended to set the FS to at least 5 to reduce analysis time without compromising the quality of the final output. Selected parameters are confirmed by clicking on the button: *“Update Settings”*. It is necessary to use identical parameters (t%, FS) in all control and treated experiments to ensure precise results.

The analysis process starts by clicking on the *“Analyze”* button and the final data – black and white (or red-original frame background, depending on the script) PNG image will be saved in the selected folder. The whole process took around 35-40 seconds for the analysis of 1 minute long video file recorded at 60 fps with FS set to 5.

## RESULTS

The Graphical analysis (GA), and the Pixel Activity Analysis (PAA) could be used sequentially as shown here (Fig. 1) and described in Methods.

**Fig. 1:**
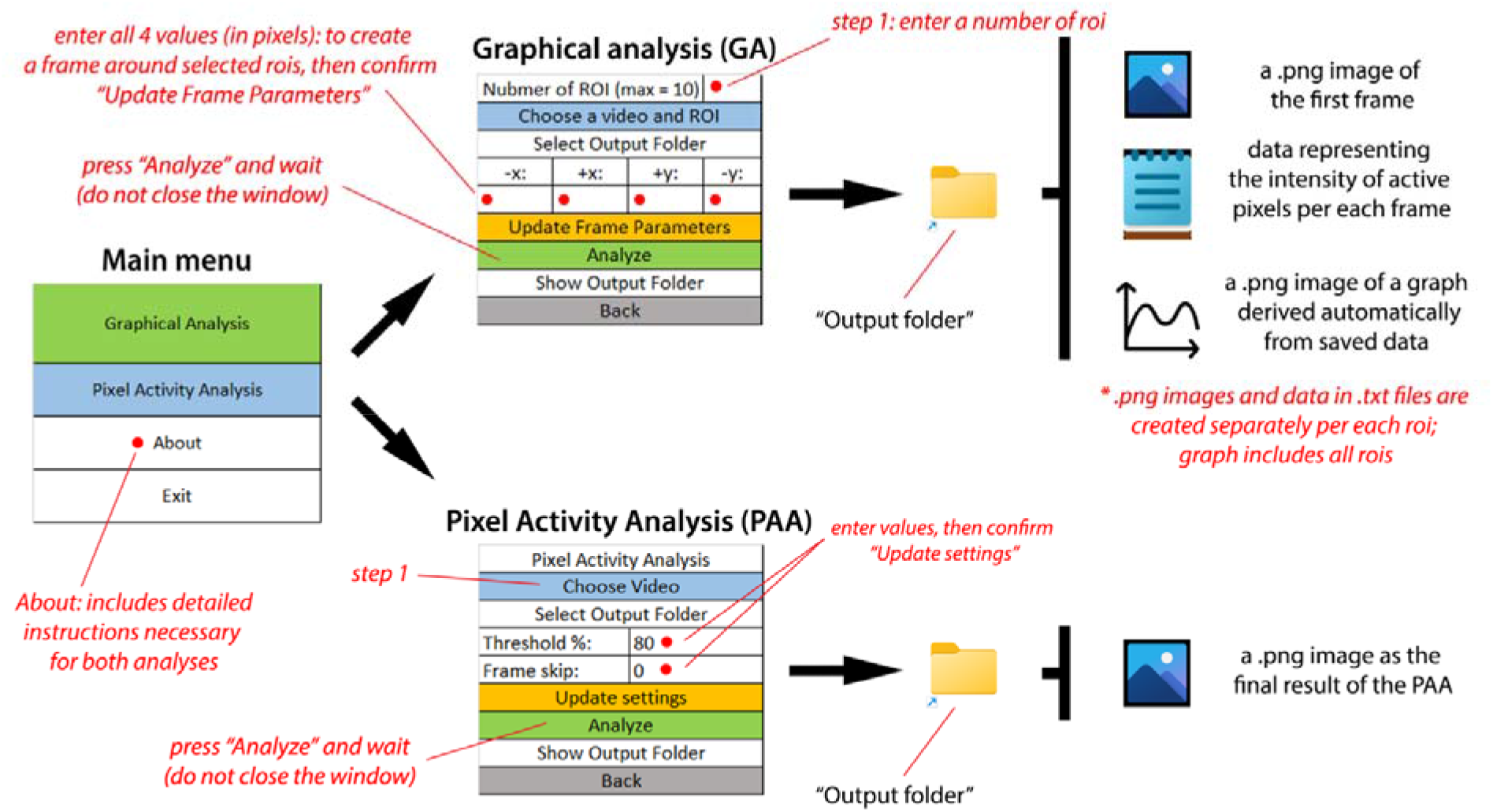
Schematic representation of TickMotion workflow. TickMotion can process video files and provides two distinct analyses: Graphical Analysis (GA) and Pixel Activity Analysis (PAA). In GA, the initial step involves entering the number of regions of interest (ROIs). Following this, the process is similar for both analyses: choosing a video file, selecting the output folder, adjusting parameters, confirming the settings and finally starting the analysis. The results are then saved in the designated folders and can be easily accessed directly from the application.

Male ticks (n=5) were dissected in physiological solution by carefully removing the dorsal cuticle and gut to expose the accessory glands (Fig. 4B). To detect muscle activity, *vas deferens* within the male accessory gland was selected as the target area (Fig. 4B, arrows; see Suppl. Fig. S1A-D). Samples were evaluated at three experimental conditions: baseline (control), after application of a muscle stimulator, and after application of a muscle inhibitor, using the GA (Fig. 2). A significant increase in signal intensity (indicating “pixel activity” uptake) was observed following stimulator application (Suppl. Video 2), compared to the lower activity levels recorded under control conditions (31.22%; Suppl. Video 1) and after inhibitor application (19.25%; Suppl. Video 3) (Fig. 2A). At the control and inhibitor stages, only weak background signals were present in the targeted region, with no distinct amplitude peaks (Fig. 2B; Suppl. Videos 1 and 3). In contrast, during stimulation, repetitive amplitude peaks (consistent with rhythmic muscle contractions) emerged after an initial short delay (~45 seconds) and occurred approximately every 3 seconds thereafter, continuing until the end of the recording (manually stopped) (Fig. 2B; Suppl. Video 2).

**Fig. 2:**
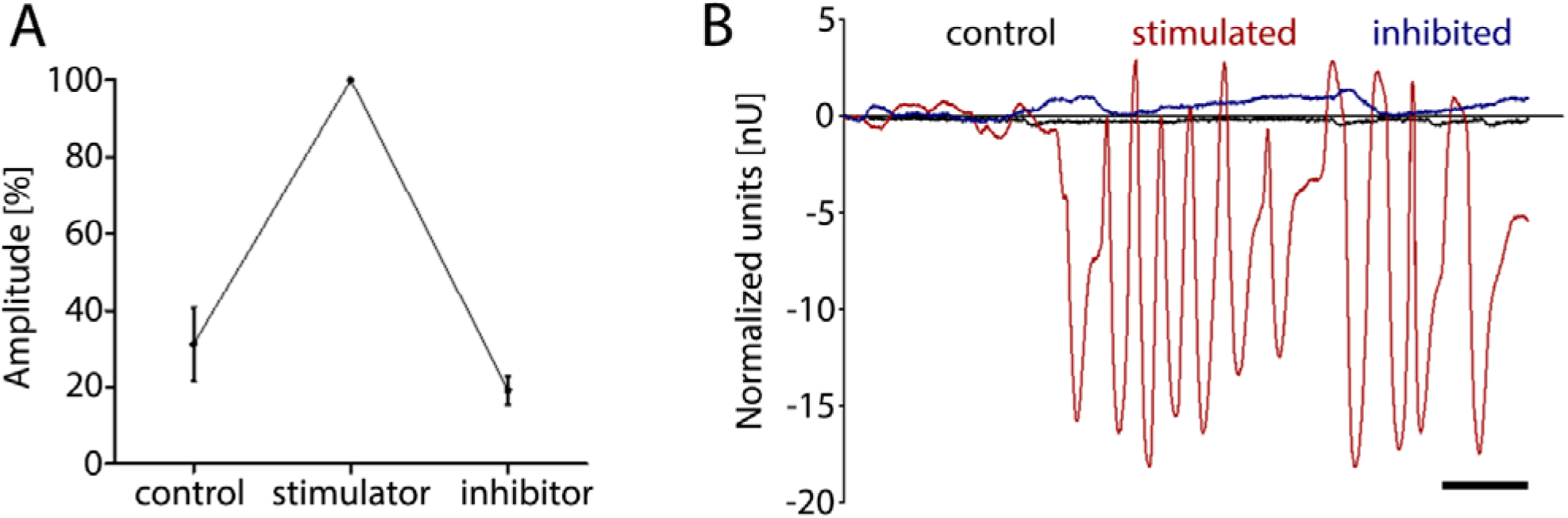
Quantification of pixel activity within selected regions of interest (ROIs) in the male accessory gland under three experimental conditions: control, stimulated, and inhibited, using the Graphical Analysis (GA). Pixel activity was measured within a defined frame area (parameters: –x = 50; +x = 50; –y = 30; +y = 30) surrounding the ROI. **(A)** Graphical representation of pixel activity differences among experimental conditions (n = 5), expressed as the percentage change in the interquartile range (IQR, Q3-Q1). **(B)** Activity progression curves over a 2-minute videos, showing temporal dynamics of pixel activity in the stimulated (red), control (black), and inhibited (blue) accessory glands (scale bar = 15 seconds).

The dorsal cuticle of partially-fed female ticks were dissected to expose the gut (Fig. 4A, arrow). Muscle contractions were stimulated (Suppl. Video 4) and analyzed using PAA with varying threshold percentage parameters: t%80, t%85, t%90, and t%95. The results demonstrated that higher t% values correspond to lower number of pixels appearing as white and indicating increased activity (Fig. 3).

**Fig. 3:**
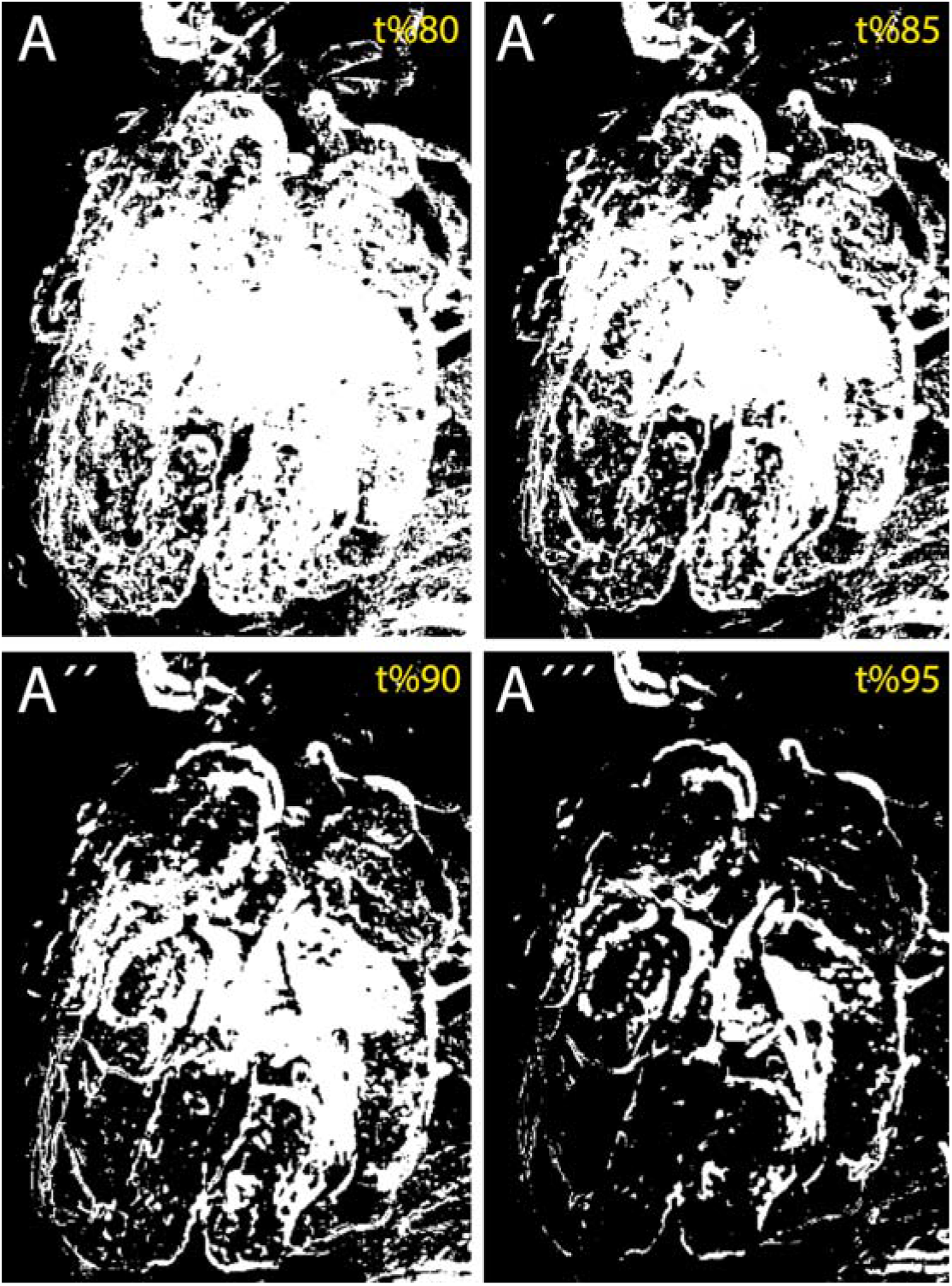
Application of different threshold percentage (t%; t%80 – t%95) on a video with contracting tick’s gut. Note, when lower t% is applied **(A)**, more active pixels (white) are highlighted in the result, comparing to higher t% **(A’-A’’’)**.

Optimizing key parameter values (i.e., t% and FS) in PAA is crucial for further functional analysis of recordings from specific time points. The highest “pixel activity” area in selected recordings following muscle stimulation was observed in the midgut of partially-fed females (Fig. 4A, arrow; Suppl. Fig. S1E) and *vas deferens* of the male accessory gland (4B, arrows; Suppl. Fig. S1A-D). Increased signal was observed after stimulation (Fig. 4A’’, 4B’’; Suppl. Videos 2 and 4) when compared with controls (Fig. 4A’, 4B’; Suppl. Videos 5 and 1).

**Fig. 4:**
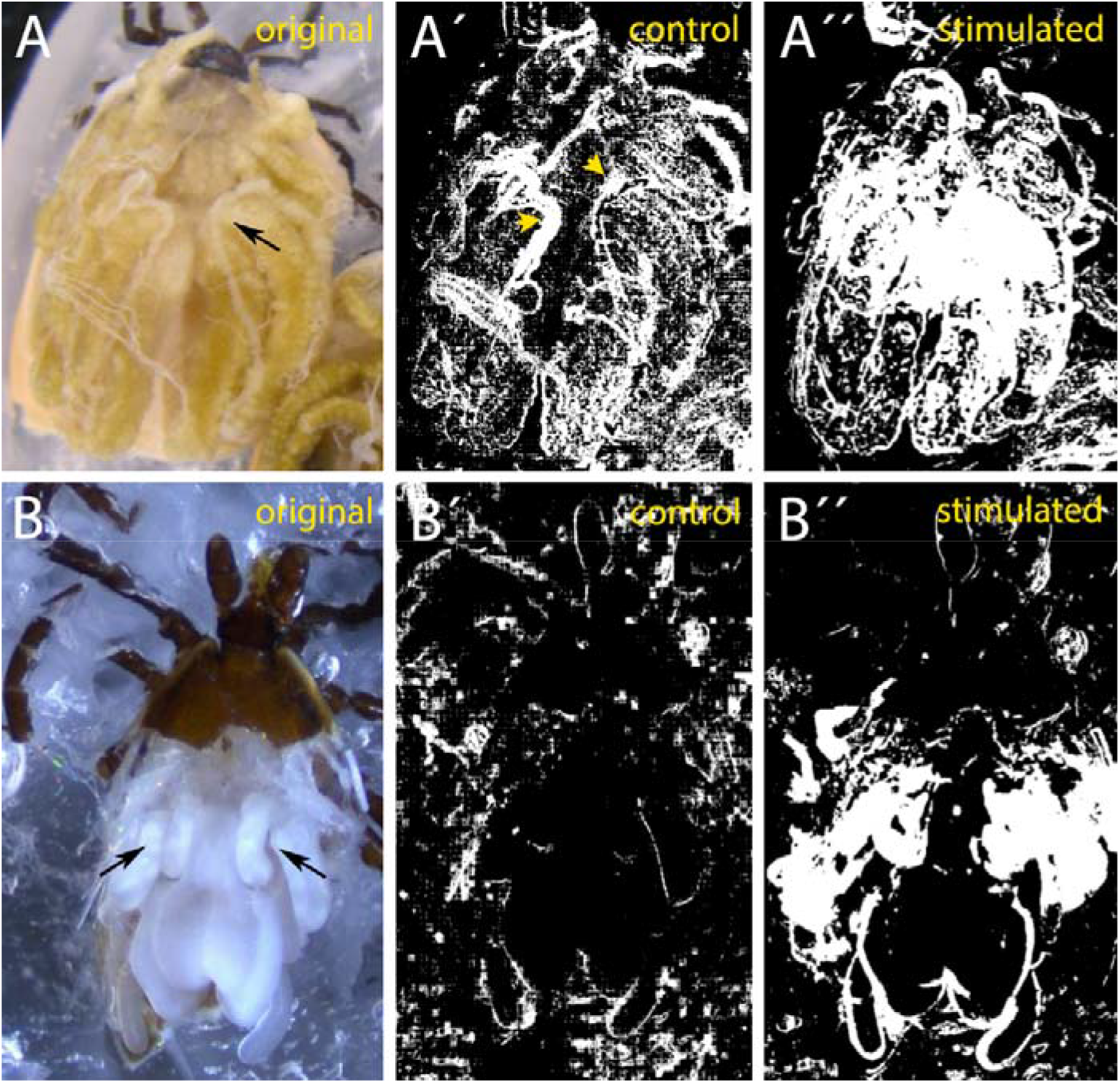
Detection of organ-specific pixel activity using TickMotion Pixel Activity Analysis (PAA) with parameters: t%=85, FS=5. **(A, B)** Visualization of muscle activity in the female midgut (A-A’’, arrow) and male *vas deferens* (B-B’’; arrows) under control (A’, B’) and stimulated (A’’, B’’) conditions. In the control female midgut (A’), the only prominent white regions (yellow arrowheads) correspond to the Malpighian tubules, which exhibited spontaneous contractile activity in all *in vitro* preparations and are considered background signal.

Second version of the script, TickMotion v_2, featuring modified PAA, was utilized for both *in vitro* detection of muscle contractions in tick tissues and *in vivo* analysis of tick movements. TickMotion v_2 produced the identical pattern in “pixel activity” (Fig. 5A, B) using the same recordings and PAA parameters as previously described (Fig. 4A’’, 4B’’) (Suppl. Videos 2 and 4). Additionally, this version was successfully applied to track a male tick’s walking on a Petri dish (Fig. 5C; Suppl. Video 6) showing the applicability of the PAA for tracking arthropod movements.

**Fig. 5:**
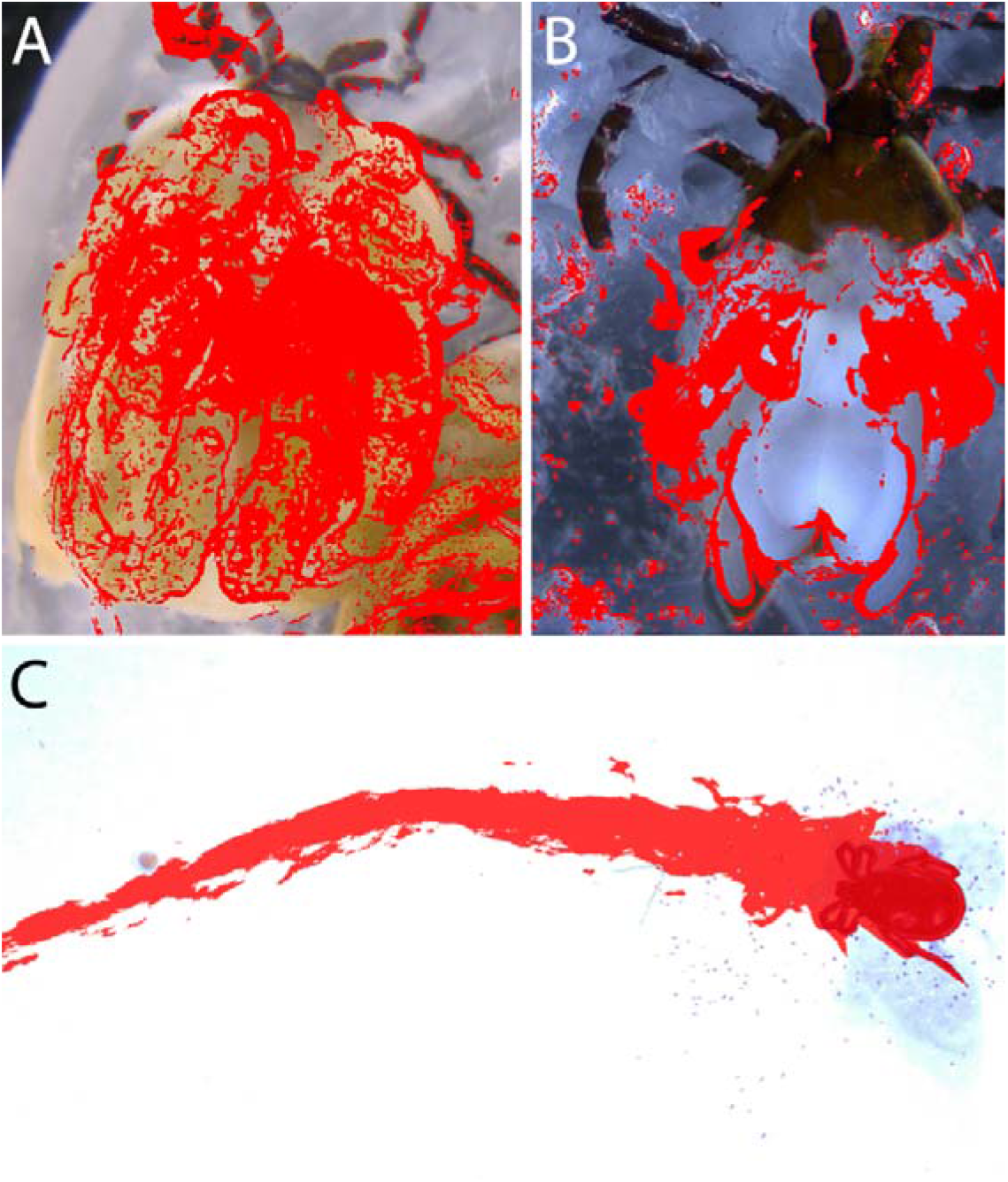
TickMotion v_2 script contains modified Pixel Activity Analysis (PAA) that enables quantification of both tissue contractions and broader movements as demonstrated in *I. ricinus*. **(A, B)** Detection of muscle contractions in a stimulated female gut (A) and male accessory gland (B), using the same PAA parameters as in Fig. 4 (t%=85, FS=5). **(C)** Analysis of whole-body movement during male tick locomotion from a 22-second video, using adjusted PAA parameters: t%=90, FS=0.

## DISCUSSION

We present here a new Python-based tool, TickMotion, developed for post hoc analysis of muscle contractions and movements from video recordings in insects and other arthropods. We successfully tested the application on *I. ricinus* both *in vivo* by tracking male tick walking, and *in vitro* by analyzing muscle contractions in the male accessory glands and the midgut of partially-fed females. Our results demonstrate that TickMotion can distinguish specific patterns (i.e., frequency and amplitude of muscle contractions) between control and treated samples. Moreover, the combined use of Graphical Analysis (GA) and Pixel Activity Analysis (PAA) provides comprehensive insights through both qualitative and quantitative assessments.

The GitHub repository hosts two versions of the script: “TickMotion” and “TickMotion v_2”, with the latter serving as an upgraded version of the original. The second version was specifically tested in experiments tracking male tick walking and demonstrated notable improvements. Unlike the original script, which primarily provides black-and-white imaging, “TickMotion v_2” preserves detailed visual information, including tissue structures, shapes, and colors, which are derived from the first video frame. For example, when studying muscle contractions in specific part of tissue or organs, or movement in particular directions, the ability to observe color variations and structural details provides valuable insights that might be lost in a binary or grayscale representation. As a result, “TickMotion v_2” script offers greater flexibility and precision, making it especially beneficial for experiments requiring clear differentiation between moving structures and their surroundings.

Several established tools, such as EthoVision XT, DeepLabCut, and ANY-Maze, are widely used for analyzing animal behavior, including tracking, pose estimation, and movement analysis in larger animals. While these programs are powerful and comprehensive, they are often too complex, resource-intensive, and expensive for routine use in arthropod muscle contraction and fine-scale movement studies. For example, EthoVision XT is primarily designed for rodents and small laboratory animals, and DeepLabCut and ANY-Maze, though effective for behavioral assays, lack specialized features tailored to muscle contraction analysis in small arthropod tissues (Bühler et al., 2023; Mathis et al., 2018; Bailoo et al., 2010). Therefore, there remains a need for a simpler, more accessible tool specifically optimized for arthropod physiology research.

TickMotion stands out due to its user-friendly interface and streamlined software design, which facilitate efficient and intuitive video analysis. Moreover, TickMotion is freely available on GitHub, easy to install, and exhibits relatively high sensitivity during measurement. Finally, its customizable parameters using GA and PAA – enhance result accuracy and reliability, making it an accessible yet effective tool for arthropod movement studies.

It is important to note that TickMotion analyzes the intensity of pixel activity in a single video layer. This means it cannot selectively isolate a tissue or segments at specific depths within the video all pixels within the selected area are analyzed indiscriminately. Consequently, the output results may be affected by any “contamination” in the video, e.g., shadow, unspecific movement, or anything that will disturb the recording. Despite these limitations, the analysis provides highly precise data. During the examination of muscle dynamics in the male accessory gland using GA, a small number of frames were manually excluded due to external disturbances, such as brief shaking artifacts – 4 out of approximately 8,500 frames after inhibition and 1 out of 5,000 frames after stimulation. These frames were removed not because they affected the data analysis based on the interquartile range (IQR), but to improve the visual clarity and progression of the resulting curve, as the values were abnormally high or low and would have distorted the overall trend. This manual exclusion ensured measurement accuracy without compromising overall data integrity. All video analyses were performed on a computer running on a Windows OS, 13th Gen Intel(R) Core(TM) i7-1355U, 1.70 GHz, 32 GB RAM, 64 bit. Using a computer or laptop with different specifications may negatively impact the analysis process.

## ACKNOWLEDGEMENT

This work was supported by the Slovak grant agencies: APVV-21-0431, APVV-23-0168, VEGA-2/0070/23, VEGA-2/0037/23, and TUBITAK-2024-01. Matej Medla is a recipient of Štefan Schwartz support fund 2025/OV2/018.

## REFERENCES

Bailoo JD, Bohlen MO, Wahlsten D. The precision of video and photocell tracking systems and the elimination of tracking errors with infrared backlighting. J Neurosci Methods. 2010; 188(1):45–52. doi:10.1016/j.jneumeth.2010.01.035

Bühler D, Power Guerra N, Müller L, Wolkenhauer O, Düffer M, Vollmar B, Kulha A, Wolfien M. Leptin deficiency-caused behavioral change – a comparative analysis using EthoVision and DeepLabCut. Front Neurosci. 2023; 17:1052079. doi:10.3389/fnins.2023.1052079

Hermanson JW, Altenbach JS. Functional anatomy of the primary downstroke muscles in the pallid bat. J Mammal. 1981; 62:795–800. doi:10.2307/1380600

Hewes RS, Snowdeal EC, Saitoe M, Taghert PH. Functional redundancy of FMRFamide-related peptides at the Drosophila larval neuromuscular junction. J Neurosci. 1998; 18:7138–7151.

Hu Z, Pym EC, Babu K, Vashlishan Murray AB, Kaplan JM. A neuropeptide-mediated stretch response links muscle contraction to changes in neurotransmitter release. Neuron. 2011; 71(1):92–102. doi:10.1016/j.neuron.2011.04.021

Kim DH, Han MR, Lee G, Lee SS, Kim YJ, Adams ME. Rescheduling behavioral subunits of a fixed action pattern by genetic manipulation of peptidergic signaling. PLoS Genet. 2015; 11(9):e1005513. doi:10.1371/journal.pgen.1005513

Mathis A, Mamidanna P, Cury KM, Abe T, Murthy VN, Mathis MW, Bethge M. DeepLabCut: markerless pose estimation of user-defined body parts with deep learning. Nat Neurosci. 2018; 21(9):1281–1289. doi:10.1038/s41593-018-0209-y

Matsumoto S, Kutsuna N, Daubnerová I, Roller L, Žitňan D, Nagasawa H, Nagata S. Enteroendocrine peptides regulate feeding behavior via controlling intestinal contraction of the silkworm Bombyx mori. PLoS One. 2019; 14(7):e0219050. doi:10.1371/journal.pone.0219050

Onken H, Moffett SB, Moffett DF. The anterior stomach of larval mosquitoes (Aedes aegypti): effects of neuropeptides on transepithelial ion transport and muscular motility. J Exp Biol. 2004; 207:3731–3739.

Pflüger HJ, Duch C. Dynamic neural control of insect muscle metabolism related to motor behavior. Physiology (Bethesda). 2011; 26(4):293–303. doi:10.1152/physiol.00002.2011

Šimo L, Park Y. Neuropeptidergic control of the hindgut in the black-legged tick Ixodes scapularis. Int J Parasitol. 2014; 44(11):819–826. doi:10.1016/j.ijpara.2014.06.007

Vassallo M, Pérez-Eid C. Comparative behavior of different life-cycle stages of Ixodes ricinus (Acari: Ixodidae) to human-produced stimuli. J Med Entomol. 2002; 39(1):234–6. doi: 10.1603/0022-2585-39.1.234.

Vo-Doan TT, Dung VT, Sato H. A cyborg insect reveals a function of a muscle in free flight. Cyborg Bionic Syst. 2022; 2022:9780504. doi:10.34133/2022/9780504

Whitley E, Ball J. Statistics review 1: presenting and summarising data. Crit Care. 2002; 6(1):66–71. doi:10.1186/cc1455

Yoshida S, Takaki K, Yamawaki Y. Roles of muscle activities in foreleg movements during predatory strike of the mantis. J Insect Physiol. 2023; 145:104474. doi:10.1016/j.jinsphys.2022.104474

